# INFLUENCE OF LIFESTYLE ON BRAIN SENSITIVITY TO CIRCULATING INSULIN-LIKE GROWTH FACTOR 1

**DOI:** 10.1101/2025.02.27.640579

**Authors:** J. Zegarra-Valdivia, M.Z. Khan, A. Putzolu, R. Cipriani, J. Pignatelli, I. Torres Aleman

## Abstract

Life style conditions such as social relationships and diet impinge on mood homeostasis, a mechanism that becomes dysregulated in high-incidence mental illnesses such as depression or Alzheimer’s dementia (AD). Since insulin-like growth factor 1 (IGF-1) modulates mood and its blood levels are altered both in AD and in affective disorders, we investigated whether its activity was altered in the brain of mice submitted to isolation or fed with a high-fat diet (HFD). As in humans, both life style conditions increased anxiety and depression-like behavior. Significantly, both life style conditions abrogated neuronal responses to systemic IGF-1. Thus, enhanced neuronal activity in response to intraperitoneal IGF-1, as determined by Ca^++^ fiber-photometry in the prefrontal cortex, was lost in isolated or HFD-fed mice. However, only the latter had elevated serum IGF-1 levels. These findings suggest that loss of brain IGF-1 input may contribute to mood disturbances observed in lonely and obese subjects. Furthermore, they provide additional insight into the heightened risk of depression and Alzheimer’s disease associated with these conditions. Importantly, since the reduction of IGF-1 activity in the brain is not consistently mirrored by its serum levels, serum measurements do not reliably reflect brain IGF-1 activity.

## Introduction

Life style influences brain health ^1^. We previously postulated that IGF-1 input to the brain may help explain the effect of life-style factors on brain health as conditions such as stress or inflammation, associated to life style factors such as sociality and diet, respectively, interfere with IGF-1 activity ^2^. Indeed, IGF-1 is a wide-spectrum neuromodulator, highly abundant in the circulation ^3^. Circulating IGF-1 crosses the blood-brain-barrier ^4^, and exerts important functions at central level, affecting energy balance ^5^, mood ^6^, cognition ^7^, and sociability ^8^. Low serum IGF-1 levels increase vulnerability to stress ^9^, and cause depression ^10^, whereas obese ^11^ or depressed ^12^ individuals show either high bioactive IGF-1 or high serum IGF-1 levels, respectively, hinting to a status of IGF-1 resistance ^13^. Recently, we found in mice that both stress ^2^, and overweight ^14^ markedly reduce the entrance of IGF-1 into the brain, pointing to an underlying brain IGF-1 resistance probably at the blood-brain-barrier (BBB).

We now analyzed the effect of isolated housing or high-fat diet (HFD) in adult mice of both sexes on mood and the effect of systemic IGF-1 in their brains measured by neuronal activity. Both life-style conditions worsened mood and were accompanied by a lack of neuronal activation by systemic IGF-1 injection. However, only HFD-fed mice showed elevated serum IGF-1. These results emphasize the need to develop functional assays to determine the activity of IGF-1 in target tissues such as the brain to better understand the role of this growth factor in health and disease.

## Materials and Methods

### Animals

Adult male and female C57BL/6J mice (Janvier Labs, France) were used in a balanced manner. Mice were kept in a room with controlled temperature (22°C) under a 12-12h light-dark cycle, and water ad libitum. Mice were fed with a control or high fat diet (HFD) containing 45% fat in the diet + 1.25% cholesterol (ref E15744-34, sniff Spezialdiäten GmbH; Germany). All experimental protocols were performed during the light cycle between 12,00 – 17,00 hours. Animal procedures followed European guidelines (2010/63/EU) and were approved by the local Bioethics Committee (M20-2022-009 PIBA).

### Experimental design

Mice were submitted to isolated (1 animal/cage) or standard group housing conditions (4-5/cage), and fed with standard or HFD diet for 16 weeks (Suppl Fig 1A). Both male and female HFD-fed mice started to be overweight at around 3-4 weeks (Suppl Fig 1B). HFD mice show glucose intolerance along with hyperinsulinemia and insulin resistance after 10 weeks, as already published elsewhere ^15^. Various tests were used to confirm anxiety/depression, and serum corticosterone was also measured. A sexual dimorphism in feeding behavior was observed, (Suppl Figure 2A), and, as expected, females gain less weight than males (Suppl Figure 1B), suggesting that their energy expenditure is higher. At the end of the experimental period, blood and brain tissue were collected from each animal.

### Behavioral tests

Both males and females were analyzed together as no sex differences in behavior were appreciated. After each trial, apparatuses were cleaned with 70% ethanol to remove scent cues. Mice were acclimated to the testing room for 30 minutes before the test, which was conducted under dim lighting.

#### Zero maze

To assess anxiety-like behavior in mice we evaluated their preference for open versus closed sections of an elevated, circular track. The maze consists of two open and two closed quadrants, elevated 40–60 cm above the floor, with each quadrant spanning 90°. Each mouse was placed at the border between an open and a closed section, facing the open area, and allowed to explore for 5–10 minutes while being video recorded. Time spent in open and closed sections was determined, calculating the percentage of time spent in open sections and the number of entries, with increased time in open sections indicating lower anxiety-like behavior and decreased time suggesting higher anxiety.

#### Forced swim test

To assess depressive-like behavior, we used the forced swim test as described before ^16^. Mice were placed in a glass cylinder (12 cm diameter, 29 cm height) filled with water (23°C) to a height of 15 cm (to avoid climbing). The animals keep swimming until they give up, or they alternate between swimming and floating. A 6 min test session was video recorded and the last 4 min were scored (immobility time). *Y-maze:* Working memory was assessed by recording spontaneous activity using a Y-maze, as reported ^17^. The maze was made of grey acrilic plastic, and each arm was 25 cm long, 14 cm high, 5 cm wide, and positioned at equal angles. After each 8 minutes trial, the maze was cleaned with 70% ethanol to remove any olfactory cues. An off-line analysis of the videos was carried out to obtain the sequence of entries during the test. Alternate behavior was calculated as the percentage of real alternations (number of triplets with non-repeated entries) versus total alternation opportunities (total number of triplets).

#### Social behavior

We studied social novelty/preference in control mice, as described by others ^18^. Each mouse was placed in a cage with three compartments (one central and two lateral arms); in each compartment, we added a grid with one stranger mouse or an empty grid to assess social affiliation (intention to stay with the same species). We leave the mouse to explore for 10 minutes and record the time of direct interaction. Then, we cleaned the three chambers with ethanol and placed the mouse again in the center chamber. We include the previous stranger mouse in the same arm (now named “familiar mouse”). In the empty space we include a new mouse (“stranger mouse”) and leave the animal free to explore and record the time of direct interaction.

### Ca^++^ Fiber-photometry

Fiber-photometry was used to assess in vivo neuronal activity through monitoring calcium dynamics, which serve as a proxy for neuronal firing. We stereotaxically injected an AAV-GCaMP virus (ssAAV-9/2-hSyn1-jGCaMP8m-WPRE-SV40p(A); Viral Vector Facility, ETH, Switzerland, 6.4 × 10E^12^ vg/ml, 500 nl unilaterally injected) encoding a calcium-sensitive fluorescent indicator into the prefrontal cortex (AP=1.7; ML=0.5; DV=±1.5) four weeks before the experiments, at a rate of 100 nl/min and the Hamilton syringe was withdrawn 10 min later. Mice were anaesthetized with isoflurane (Zoetis, USA) administered with a nose mask (David Kopf Instruments, France), and placed on a stereotaxic frame (Stoelting Co, USA) on a heating pad and tape in their eyes to protect them from light.

#### Peak Detection Analysis

In the day of the experiment, mice received a bolus injection of IGF-1 dissolved in saline and intraperitoneally (ip) injected (1µg/g body weight), or fasted for 20-24 h and ip injected with glucose (2 grs/kg). We use the R811 Dual Color Multichannel Fiber Photometry System (RWD Life Science, China) to enable real-time detection and quantification of transient fluorescence signals (ΔF/F or Z-score) associated with neuronal activity in freely moving animal. In the preprocessing pipeline, 470 nm was selected as the signal wavelength, while 410 nm was designated as the control channel to correct movement artifacts and non-specific fluctuations. A smoothing filter (W coefficient = 15) was applied to reduce high-frequency noise without distorting signal integrity. To correct for fluorescence bleaching effects, baseline correction (PLS Fit algorithm, β coefficient = 8) was implemented, which is particularly suited for short and regular cycles of fluorescence decay. Furthermore, motion correction was applied to remove confounding artifacts caused by fiber displacement, ensuring that detected fluorescence variations represented genuine neuronal activity.

Following preprocessing, Peak Statistics parameters were configured for signal quantification. To refine the robustness of peak identification, the Median Absolute Deviation (MAD) Duration was set to 0.00–5.00 seconds, enabling precise calculation of the MAD value as a reference threshold. The Threshold for peak detection was adjusted to 1.00 MAD, ensuring that only fluorescence changes surpassing this criterion were considered significant. Additionally, Peak distances were set to 5.0 seconds, allowing for selecting the highest peak within each time frame while disregarding redundant peaks within the specified interval.

The Statistics Duration was set to 0–300 seconds at basal, defining the analysis window for peak detection, to assess neuronal activity changes, during which the peak count, frequency (Hz), and mean peak amplitude (% ΔF/F or Z-score) were quantified, with baseline activity considered the 100% reference. Subsequently, following the administration of IGF-1 or glucose, peak activity was evaluated within 5 min intervals, allowing for comparative analysis of neuronal responses to these interventions.

### ELISA

Serum IGF-1 (Biotechne R&D Systems, USA), brain pSer^318^IRS-1/IRS-1 (Thermo-Fisher, USA), and serum corticosterone (Enzo Life Sciences, Australia) levels were determined using commercial ELISAs, following the manufacturer’s instructions. Blood was collected at the same time of the day in all experimental groups to minimize circadian variations.

### Statistics

Statistical analysis was performed using Graph Pad Prism 10 software (USA). Depending on the number of independent variables, normally distributed data (Kolmogorov-Smirnov normality test), and the experimental groups compared, we used either one-way ANOVA or two-way ANOVAs by Tukey’s or Sidak’s multiple comparison test, respectively. The sample size for each experiment was chosen based on previous experience and aimed to detect at least a p<0.05 in the different tests applied, considering a reduced use of animals. Results are shown as mean ± standard error (SEM) and p values coded as follows: *p< 0.05, **p< 0.01, ***p< 0.001.

## Results

### Life-style modulates mood

Since changes in mood elicited by isolation or HFD were sex-independent (Suppl Figure 2B,C), we include the results of males and females together. Both life-style conditions elicited higher anxiety, as determined by the Zero maze (Figure 1A).

**Figure 1:**
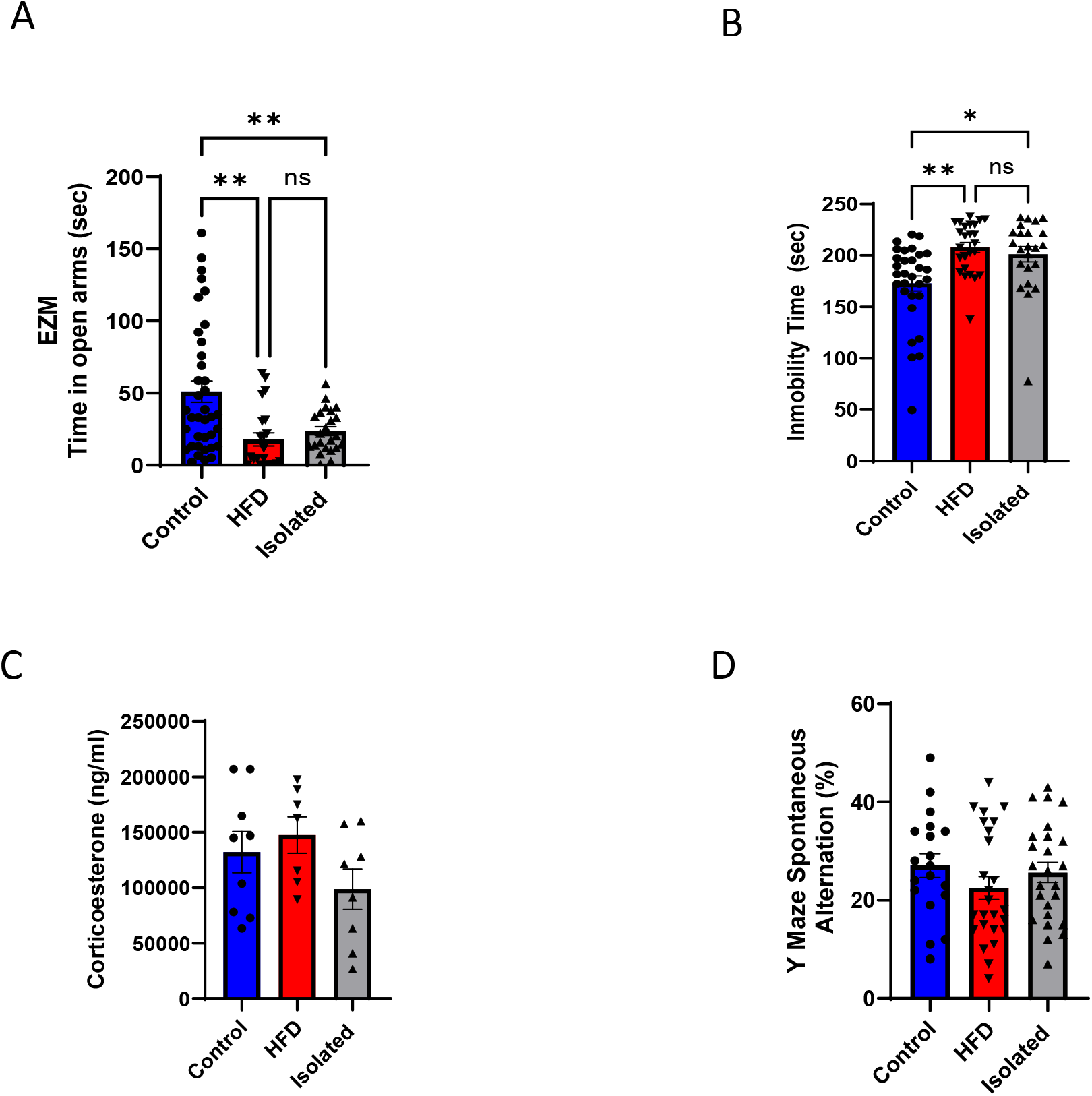
Effects of life-style on mood. **A**, Anxiety levels in the different groups was measured in the Zero maze. Both isolated and HFD-fed mice showed higher anxiety, as determined by less time spent in the open arms of the maze; Controls (n=37), HFD (n=22), Isolated (n=23). **B**, Depression-like behavior was determined using the Forced Swim Test (FST). Both isolated and HFD-fed mice show increased depressive symptoms; Controls (n=30), HFD (n=25), Isolated (n=23). **C**, No significant changes were found in serum corticosterone levels in isolated and HFD-fed mice, compared to controls; Controls (n=9), HFD (n=7), Isolated (n=8). **D**, Spatial memory was assessed measuring alternation behavior in the Y-maze test. No changes in this behavior were seen as compared to the control group; Controls (n= 19), HFD (n=25), Isolated (n=25). P value is as follow: *p<0,05; **p<0,01; ***p<0,001 in this and following figures.

Regarding depressive-like behavior determined by the forced swim test (FST), we observed that both experimental groups spent more time floating, indicating a depression-like behavior (Figure 1B). Serum corticosterone levels in isolated mice were non-significantly lower than in control mice, whereas in HFD-fed mice were slightly, but not significantly, increased (Figure 1C). We then assessed cognition using the Y-maze spontaneous alternations test, which measures working memory, and neither group showed alterations compared to control mice (Figure 1D).

### Life-style modulates neuronal responses to systemic IGF-1

Next, we used fiber-photometry with viral-transduced Ca^++^ sensors to assess the effect of IGF-1 on neuronal activity in the prefrontal cortex (PFC) of the different experimental groups through monitoring Ca^++^ dynamics, which serve as a proxy for neuronal firing ^19^. The PFC was chosen because is an area involved in mood homeostasis ^20^, influenced by social stress ^21^, and by diet ^22^. Ca^++^ dynamics were monitored to determine PFC responses to a systemic (ip) bolus injection of IGF-1 (Figure 2A). We observed that the activation seen in control mice after ip IGF-1, was absent in both isolated and HFD-fed mice (Figure 2B,C). Furthermore, since IGF-1 is involved in brain glucose handling ^23^, and glucose stimulates neuronal activity ^24^, we tested whether glucose modulates neuronal activity in isolated or HFD-fed mice following the same procedure (Figure 2A), and found loss of responses to glucose too (Figure 3), which agrees with blunted brain IGF-1 activity.

**Figure 2:**
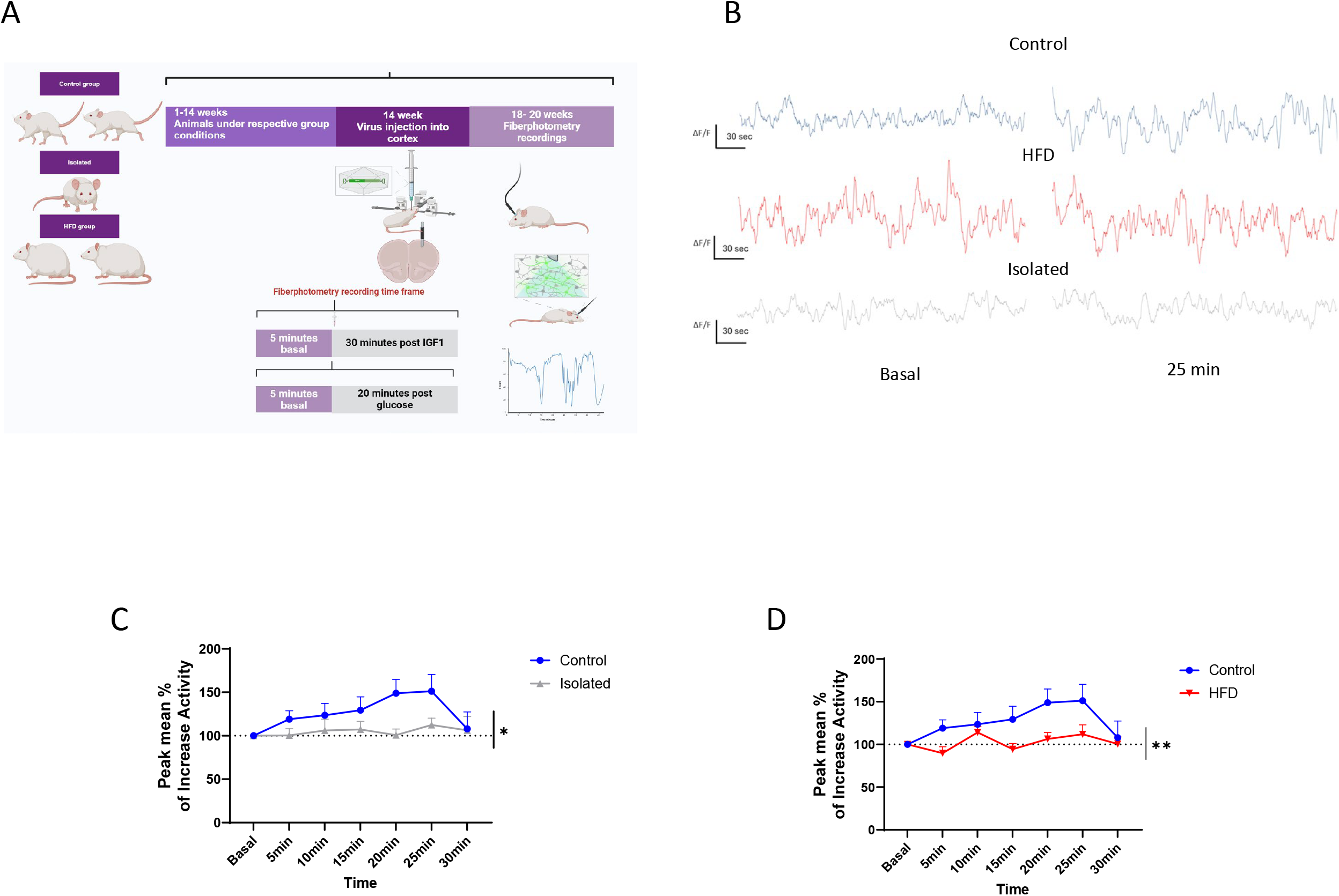
Neuronal responses to systemic IGF-1. **A**, Diagram of fiber-photometry experimental set up. **B**, Representative recordings of the three groups are shown at basal condition and at 25 min. **C**, Neuronal activity after ip IGF-1 administration was significantly increased in control mice over isolated mice. **D**, HFD-fed mice did not respond to ip IGF-1 either, as compared to controls; Controls (n= 11), HFD (n=6), Isolated (n=6).

**Figure 3:**
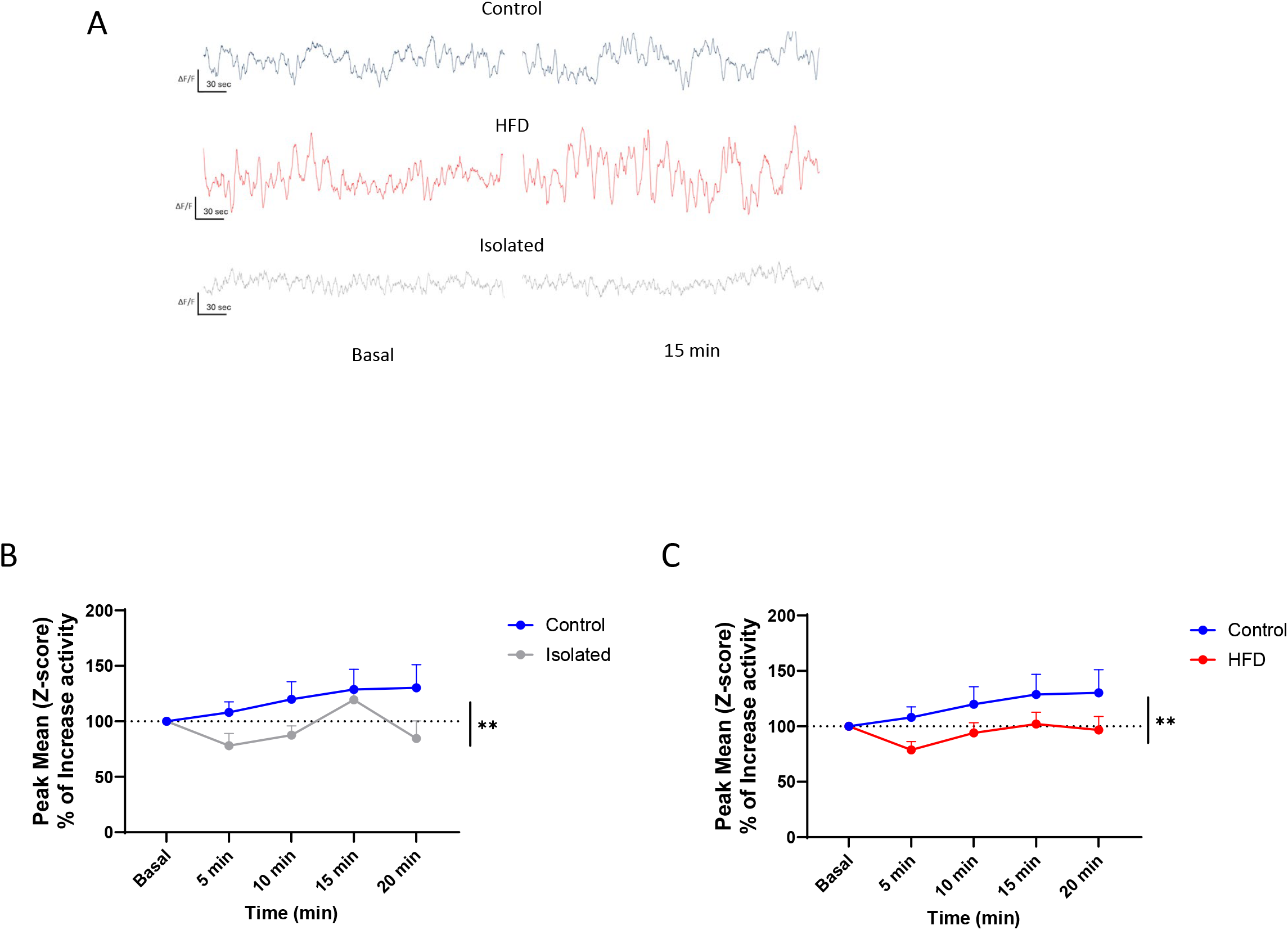
Neuronal responses to systemic Glucose. **A**, Representative fiber-photometry traces over time are displayed, showing baseline activity and recordings taken at the 15-minute mark. **B**, Neuronal activity after ip glucose was significantly increased in control mice over isolated mice. **C**, HFD-fed mice did not respond to glucose load, as compared to control mice; Controls (n=9), HFD (n=9), Isolated (n=7).

Next, loss of sensitivity to IGF-1 was determined measuring levels of phosphorylated Serine (pSer) IRS-1. This docking protein of the IGF-1 receptor modulates its sensitivity when phosphorylated in Serine residues ^25,26^. We found similar levels of cortical pSer^318^IRS-1 in the three experimental groups (Figure 4A). As already reported, we observed increased serum IGF-1 levels in HFD-fed mice ^14^, while isolated mice show no changes in serum IGF-1 (Figure 4B), even though in control mice, serum IGF-1 levels directly correlate with sociability (Figure 4C).

**Figure 4:**
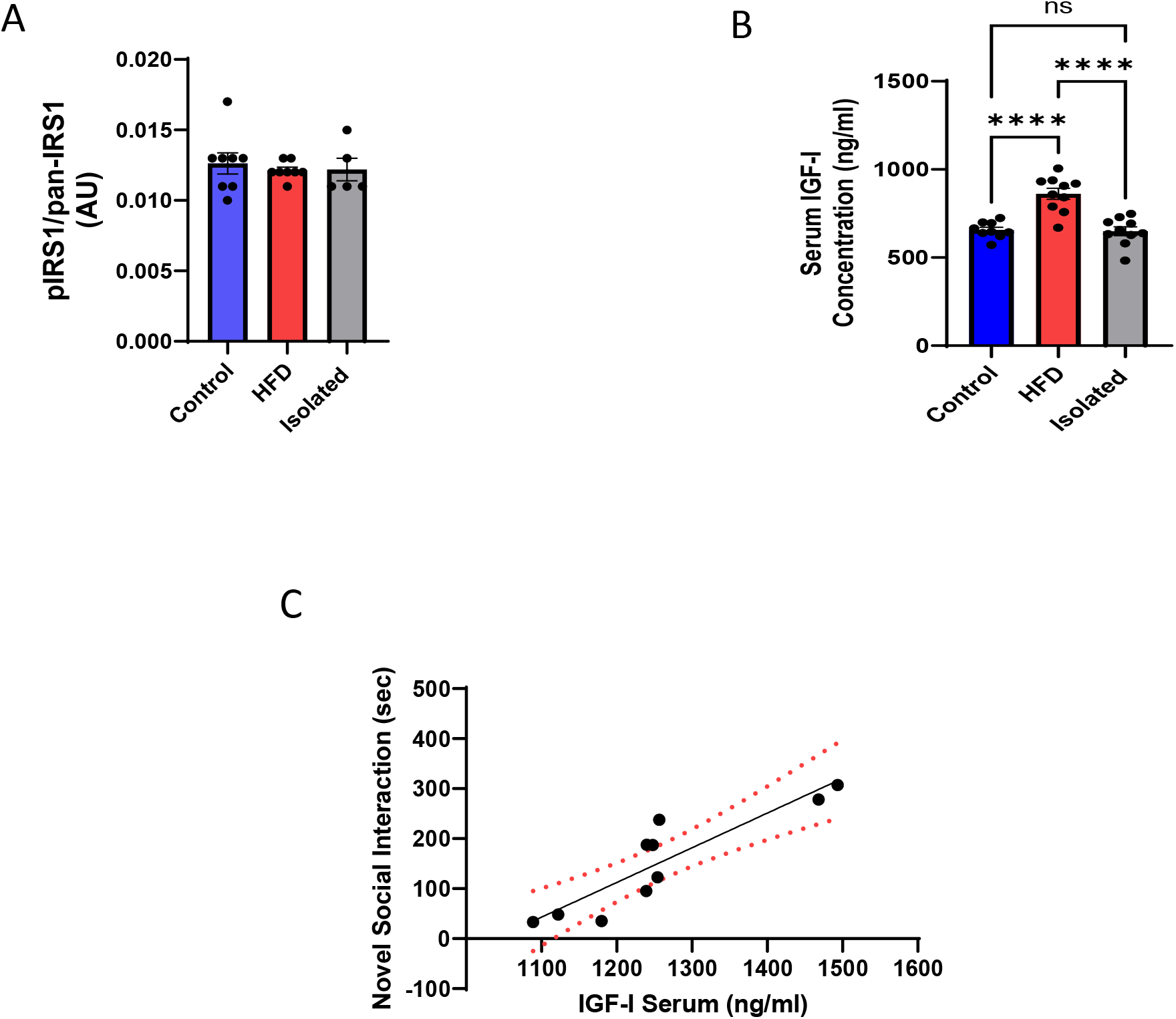
Modulation of brain IGF-1 activity by isolation and HFD. **A**, pSer^318^IRS-1 levels in cortex of the different experimental groups. Results are shown as the ratio of pSer^318^IRS1/total IRS1; Controls (n=8), HFD (n=8), Isolated (n=5). **B**, Blood IGF-1 levels increase in HFD-fed, but not in isolated mice; Controls (n=9), HFD (n=10), Isolated (n=10). **C**, Serum IGF-1 levels significantly correlate in control mice with the level of social interaction in the social novelty test. The higher the IGF-1 levels in blood, the greater the degree of sociability (n=10).

## Discussion

These results indicate that poor life style conditions such as isolated housing or imbalanced dieting are associated to a loss of activity of IGF-1 in the brain. As circulating IGF-1 access the brain ^4,27^, and modulates mood, cognition ^7,9^, and glucose handling^23^, it seems likely that reduced brain sensitivity to circulating IGF-1 input contributes to affective disturbances that are commonly seen in stressed or overweight individuals ^28,29^, and that we also observed in this mouse study. Of note, when both detrimental life-style conditions are combined, a situation that is frequent in humans, the observed mood disturbances were the same as when mice were submitted to just one condition (not shown).

Previous evidence indicates that serum IGF-1 levels are dysregulated in obese and depressive subjects ^6,30,31^. Since different processes including inflammation ^32^ or cellular senescence ^33^, are proposed to link obesity to mental illness, and multiple others (neuroinflammatory, neuroendocrine, epigenetic and metabolic) are postulated to link stress with mental illness ^34^, we consider that IGF-1 activity in the brain, which is also involved in anti-inflammatory, anti-oxidant, and metabolic mechanisms ^35^, should be considered an additional contributing factor. Indeed, neuronal activation by systemic glucose was absent in isolated or HFD mice, an indirect indication that brain IGF-1 activity is deteriorated, as this growth factor participates in brain responses to circulating glucose ^23^. As changes in IGF-1 activity have also been postulated to be involved in depression ^6^ and AD pathologies ^36^, alterations in this growth factor may help explain why loneliness and obesity are risk factors for these two psychiatric conditions ^35^.

We previously discussed that defining IGF-1 activity in the brain is not easy, although it could be a useful biomarker of brain health ^35^. For instance, in previous studies in overweight mice, we observed high serum IGF-1 levels in parallel with normal levels of brain IGF-1. At the same time, these mice showed reduced brain IGF-1 activity -measured by receptor activation in response to systemic IGF-1 ^14^. The present results corroborate these observations, and further illustrate the difficulty of defining brain IGF-1 activity based solely on IGF-1 levels ^37^. Using Ca^++^ fiber-photometry to determine neuronal responses to systemic IGF-1 we confirm that there is a loss of brain sensitivity to IGF-1 input in overweight mice and extend this observation to isolated mice. However, only overweight, but not isolated mice, show elevated serum IGF-1.

The latter may apparently contrast with our previous observation that mice with low serum IGF-1 levels show reduced sociability ^8^, or that serum IGF-1 levels directly correlate with sociability (present observations). Thus, while serum IGF-1 may affect sociability, social activity *per se* does not seem to influence serum IGF-1 levels.

The fact that cortical pSerIRS-1 levels were not modified in isolated or HFD mice suggests that loss of cortical sensitivity to systemic IGF-1 in these animals is not related to receptor desensitization. Rather, the passage of circulating IGF-1 into the brain ^4^ may be reduced, as shown previously by us ^38^, and for insulin by others ^39^.

It has been shown that serum IGF-1 levels are increased in neuronal IGF-1R knock-out mice showing reduced responses to IGF-1^40^, suggesting that loss of brain IGF-1 activity is reflected on peripheral IGF-1 levels. Differential effects of HFD and isolation on serum IGF-1 levels, despite both showing loss of brain IGF-1 activity, may be explained by the fact that while obesity is reported to elicit IGF-1 resistance in the whole body ^41^, isolation stress may affect only brain sensitivity to IGF-1, limiting its systemic impact. Alternatively, changes in serum IGF-1 during social stress due to unwanted isolation (loneliness) in mice may require longer than 3 months of exposure to eventually impinge on systemic IGF-1 levels too. Indeed, isolated mice did not show elevated serum corticosterone, suggesting that they were not stressed. Other authors have found in mice that combination of a more aggressive social stress (social intrusion) with obesity induces stronger behavioral changes ^42^, which further suggests that 3 months of isolation stress is a comparatively mild disturbance.

In summary, our results show reduced brain sensitivity to peripheral IGF-1 input in isolated and high-fat diet-fed mice, underscoring the need to develop reliable functional assays of IGF-1 activity in target organs such as the brain. This is important if we want to gain insight into the significance of this neurotrophic factor in brain health as a balance between peripheral and central IGF-1 activity has been postulated to be a key feature of its actions ^43^.

## LEGENDS TO FIGURES

**Supplementary Figure 1: A**, Experimental design: mice were divided into 3 groups (control, isolated and HFD) and kept during 9 weeks until behavioral assessments (until week 14). Thereafter they were submitted to fiber-photometry or culled for sample collection. **B**,**C**, Weight gain. Only HFD groups gain significantly more weight than controls, in both sexes. Differences in weight between males (B) and females (C) are observed, as expected. Male groups: Controls (n=10), HFD (n=13), Isolated (n=12).

Female groups: Controls (n=13), HFD (n=12), Isolated (n=10). The * symbol denotes significant differences between HFD and control or isolated mice, while the # symbol indicates significant differences between isolated and control mice, analyzed in a week-by-week comparison.

**Supplementary Figure 2: A**, Sexual dimorphism in feeding behavior: males under standard or HFD diet eat more than females. Males, but not females, maintained on HFD or isolated eat less than those on a normal diet (control group). Sample size as in supplementary Figure 1. **B**, No differences in performance were observed between males (n=20) and females (n=17) in the control group during the Zero Maze test. Similarly, no differences were found between males (n=11) and females (n=11) in the high-fat diet (HFD) group, nor between males (n=9) and females (n=14) in the isolated group. **C**, In the Forced Swim Test, no differences were detected between males and females in any of the experimental groups; control males (n=24) and females (n=15), HFD males (n=13) and females (n=12), Isolated males (n=9) and females (n=14).

## Acknowledgements

JZV and MZK are IKUR (Basque Government) fellows. This research was supported by PIBA21-33 program (Basque Government).

## Authors contributions

JZV designed and performed experiments, and wrote parts of the manuscript, MZK, AP, and RC performed experiments. ITA designed and wrote the study.

The authors declare no competing interests

## References

1 Mattson, M. P. & Arumugam, T. V. Hallmarks of Brain Aging: Adaptive and Pathological Modification by Metabolic States. Cell Metab 27, 1176–1199, doi:10.1016/j.cmet.2018.05.011 (2018).

2 Fernandez, A. M., Santi, A. & Torres Aleman, I. Insulin Peptides as Mediators of the Impact of Life Style in Alzheimer’s disease. Brain plasticity (Amsterdam, Netherlands) 4, 3–15, doi:10.3233/BPL-180071 (2018).

3 Fernandez, A. M. & Torres-Aleman, I. The many faces of insulin-like peptide signalling in the brain. Nat Rev Neurosci 13, 225–239 (2012).

4 Nishijima, T. et al. Neuronal activity drives localized blood-brain-barrier transport of serum insulin-like growth factor-I into the CNS. Neuron 67, 834–846 (2010).

5 Cheng, C. M. et al. Insulin-like growth factor 1 regulates developing brain glucose metabolism. Proc. Natl. Acad. Sci. U. S. A 97, 10236–10241 (2000).

6 Bot, M., Milaneschi, Y., Penninx, B. W. & Drent, M. L. Plasma insulin-like growth factor I levels are higher in depressive and anxiety disorders, but lower in antidepressant medication users. Psychoneuroendocrinology 68, 148–155, doi:10.1016/j.psyneuen.2016.02.028 (2016).

7 Trejo, J. L. et al. Central actions of liver-derived insulin-like growth factor I underlying its pro-cognitive effects. Mol Psychiatry 12, 1118–1128 (2007).

8 Zegarra-Valdivia, J. A., Santi, A., Fernandez de Sevilla, M. E., Nunez, A. & Torres Aleman, I. Serum Insulin-Like Growth Factor I Deficiency Associates to Alzheimer’s Disease Co-Morbidities. J Alzheimers Dis 69, 979–987, doi:10.3233/JAD-190241 (2019).

9 Santi, A., Bot, M., Aleman, A., Penninx, B. W. J. H. & Aleman, I. T. Circulating insulin-like growth factor I modulates mood and is a biomarker of vulnerability to stress: from mouse to man. Translational Psychiatry 8, 142, doi:10.1038/s41398-018-0196-5 (2018).

10 Mitschelen, M. et al. Long-term deficiency of circulating and hippocampal insulin-like growth factor I induces depressive behavior in adult mice: a potential model of geriatric depression. Neuroscience 185, 50–60 (2011).

11 Hawkes, C. P. & Grimberg, A. Insulin-Like Growth Factor-I is a Marker for the Nutritional State. Pediatr Endocrinol Rev 13, 499–511 (2015).

12 Deuschle, M. et al. Insulin-like growth factor-I (IGF-I) plasma concentrations are increased in depressed patients. Psychoneuroendocrinology 22, 493–503 (1997).

13 Jain, S., Golde, D. W., Bailey, R. & Geffner, M. E. Insulin-like growth factor-I resistance. Endocr. Rev 19, 625–646 (1998).

14 Herrero-Labrador, R. et al. Circulating Insulin-Like Growth Factor I is Involved in the Effect of High Fat Diet on Peripheral Amyloid β Clearance. Int J Mol Sci 21, doi:10.3390/ijms21249675 (2020).

15 Pignatelli, J. et al. Insulin-like Growth Factor I Couples Metabolism with Circadian Activity through Hypothalamic Orexin Neurons. Int J Mol Sci 23, doi:10.3390/ijms23094679 (2022).

16 Munive, V., Zegarra-Valdivia, J. A., Herrero-Labrador, R., Fernandez, A. M. & Aleman, I. T. Loss of the interaction between estradiol and insulin-like growth factor I in brain endothelial cells associates to changes in mood homeostasis during peri-menopause in mice. Aging (Albany NY) 11, 174–184, doi:10.18632/aging.101739 (2019).

17 Sarter, M., Bodewitz, G. & Stephens, D. N. Attenuation of scopolamine-induced impairment of spontaneous alteration behaviour by antagonist but not inverse agonist and agonist beta-carbolines. Psychopharmacology (Berl) 94, 491–495, doi:10.1007/BF00212843 (1988).

18 Kaidanovich-Beilin, O., Lipina, T., Vukobradovic, I., Roder, J. & Woodgett, J. R. Assessment of social interaction behaviors. J Vis Exp, doi:10.3791/2473 (2011).

19 Simpson, E. H. et al. Lights, fiber, action! A primer on in vivo fiber photometry. Neuron 112, 718–739, doi:10.1016/j.neuron.2023.11.016 (2024).

20 Botterill, J. J. et al. Dorsal peduncular cortex activity modulates affective behavior and fear extinction in mice. Neuropsychopharmacology 49, 993–1006, doi:10.1038/s41386-024-01795-5 (2024).

21 Wang, Z.-J. et al. Molecular and cellular mechanisms for differential effects of chronic social isolation stress in males and females. Molecular Psychiatry 27, 3056–3068, doi:10.1038/s41380-022-01574-y (2022).

22 Lowe, C. J., Reichelt, A. C. & Hall, P. A. The Prefrontal Cortex and Obesity: A Health Neuroscience Perspective. Trends Cogn Sci 23, 349–361, doi:10.1016/j.tics.2019.01.005 (2019).

23 Hernandez-Garzon, E. et al. The insulin-like growth factor I receptor regulates glucose transport by astrocytes. Glia 64, 1962–1971, doi:10.1002/glia.23035 (2016).

24 Macauley, S. L. et al. Hyperglycemia modulates extracellular amyloid-beta concentrations and neuronal activity in vivo. J Clin Invest 125, 2463–2467, doi:10.1172/JCI79742 (2015).

25 Greene, M. W., Sakaue, H., Wang, L., Alessi, D. R. & Roth, R. A. Modulation of insulin-stimulated degradation of human insulin receptor substrate-1 by Serine 312 phosphorylation. J. Biol. Chem 278, 8199–8211 (2003).

26 Talbot, K. et al. Demonstrated brain insulin resistance in Alzheimer’s disease patients is associated with IGF-1 resistance, IRS-1 dysregulation, and cognitive decline. J. Clin. Invest 122, 1316–1338 (2012).

27 Carro, E., Trejo, J. L., Busiguina, S. & Torres-Aleman, I. Circulating insulin-like growth factor I mediates the protective effects of physical exercise against brain insults of different etiology and anatomy. J. Neurosci 21, 5678–5684 (2001).

28 Hendrickx, H., McEwen, B. S. & Ouderaa, F. v. d. Metabolism, mood and cognition in aging: The importance of lifestyle and dietary intervention. Neurobiology of Aging 26, 1–5 (2005).

29 Ong, A. D., Uchino, B. N. & Wethington, E. Loneliness and Health in Older Adults: A Mini-Review and Synthesis. Gerontology 62, 443–449, doi:10.1159/000441651 (2016).

30 Frystyk, J., Brick, D. J., Gerweck, A. V., Utz, A. L. & Miller, K. K. Bioactive insulin-like growth factor-I in obesity. J Clin Endocrinol Metab 94, 3093–3097, doi:10.1210/jc.2009-0614 (2009).

31 Nam, S. Y. et al. Effect of obesity on total and free insulin-like growth factor (IGF)-1, and their relationship to IGF-binding protein (BP)-1, IGFBP-2, IGFBP-3, insulin, and growth hormone. Int J Obes Relat Metab Disord 21, 355–359, doi:10.1038/sj.ijo.0800412 (1997).

32 Lasselin, J. & Capuron, L. Chronic low-grade inflammation in metabolic disorders: relevance for behavioral symptoms. Neuroimmunomodulation 21, 95–101, doi:10.1159/000356535 (2014).

33 Ogrodnik, M. et al. Obesity-Induced Cellular Senescence Drives Anxiety and Impairs Neurogenesis. Cell Metab 29, 1061-1077.e1068, doi:10.1016/j.cmet.2018.12.008 (2019).

34 Atrooz, F., Liu, H. & Salim, S. Stress, psychiatric disorders, molecular targets, and more. Prog Mol Biol Transl Sci 167, 77–105, doi:10.1016/bs.pmbts.2019.06.006 (2019).

35 Zegarra-Valdivia, J. A., Pignatelli, J., Nuñez, A. & Torres Aleman, I. The Role of Insulin-like Growth Factor I in Mechanisms of Resilience and Vulnerability to Sporadic Alzheimer’s Disease. Int J Mol Sci 24, doi:10.3390/ijms242216440 (2023).

36 Watanabe, T. et al. Relationship between serum insulin-like growth factor-1 levels and Alzheimer’s disease and vascular dementia. J Am. Geriatr. Soc 53, 1748–1753 (2005).

37 Adams, M. M. et al. Stability of local brain levels of insulin-like growth factor-I in two well-characterized models of decreased plasma IGF-I. Growth Factors 27, 181–188, doi:10.1080/08977190902863639 (2009).

38 Dietrich, M. O. et al. Western Style Diet Impairs Entrance of Blood-Borne Insulin-like Growth Factor-1 into the Brain. Neuromolecular. Med 9, 324–330 (2007).

39 Kaiyala, K. J., Prigeon, R. L., Kahn, S. E., Woods, S. C. & Schwartz, M. W. Obesity induced by a high-fat diet is associated with reduced brain insulin transport in dogs. Diabetes 49, 1525–1533 (2000).

40 Kappeler, L. et al. Brain IGF-1 receptors control mammalian growth and lifespan through a neuroendocrine mechanism. PLoS. Biol 6, e254 (2008).

41 Imrie, H. et al. Vascular Insulin-like growth factor-1 resistance and diet-induced obesity. Endocrinology (2009).

42 Agrimi, J. et al. Obese mice exposed to psychosocial stress display cardiac and hippocampal dysfunction associated with local brain-derived neurotrophic factor depletion. EBioMedicine 47, 384–401, doi:10.1016/j.ebiom.2019.08.042 (2019).

43 Huffman, D. M. et al. Central insulin-like growth factor-1 (IGF-1) restores whole-body insulin action in a model of age-related insulin resistance and IGF-1 decline. Aging Cell 15, 181–186, doi:10.1111/acel.12415 (2016).

