## Supplementary figures and images for "INFLUENCE OF LIFESTYLE ON BRAIN SENSITIVITY TO CIRCULATING INSULIN-LIKE GROWTH FACTOR 1"

### Supplementary Figures

#### Slide 1
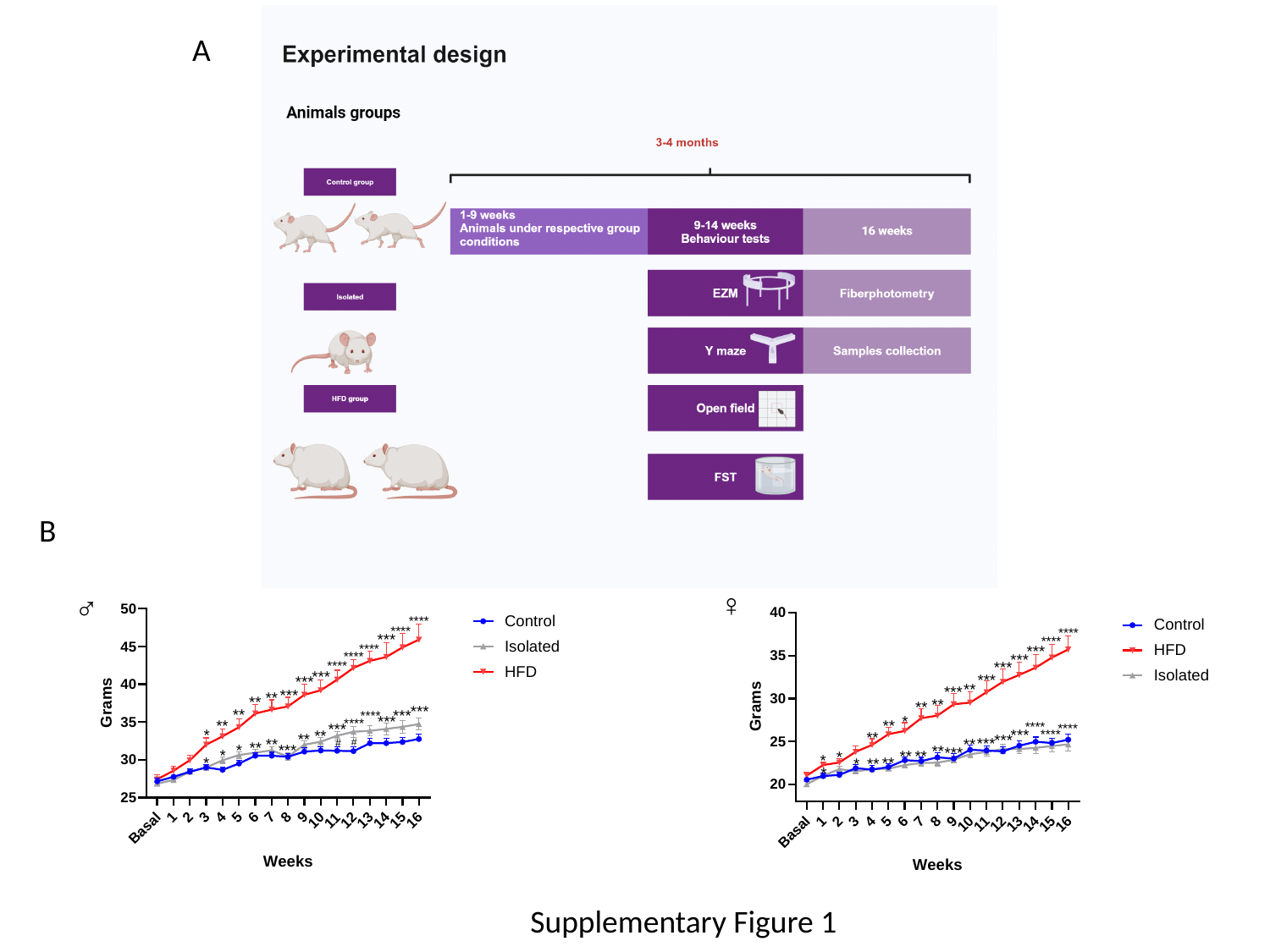

A
B
♀
♂
Supplementary Figure 1

#### Slide 2
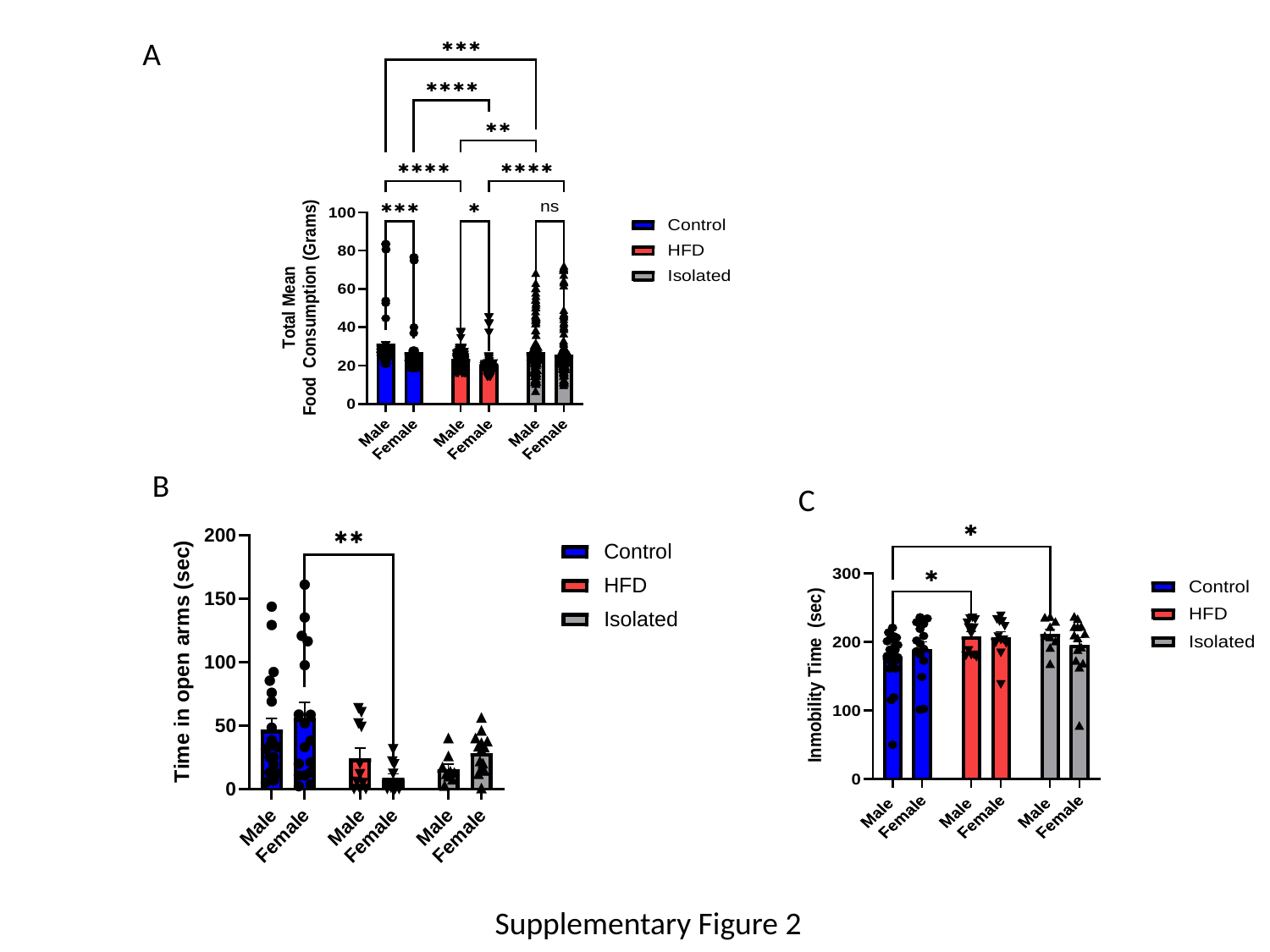

A
B
C
Supplementary Figure 2
